# Chromosomal replication origins in *Candida albicans* are genetically defined irrespective of their chromosomal context

**DOI:** 10.1101/2020.10.15.341529

**Authors:** Sreyoshi Mitra

## Abstract

Initiation of DNA replication occurs at specialized regions along chromosomes called origins. The knowledge of replication origins is imperative to understand regulation of DNA replication. The properties of replication origins in the pathogenic budding yeast *Candida albicans* remain enigmatic in the absence of the knowledge of authentic chromosomal origins. Earlier, we identified centromere proximal (pCEN) chromosomal origins on chromosomes 5 and 7 of *C. albicans*. Here, we identify another centromere-distal (dCEN) chromosomal origin by two-dimensional agarose gel electrophoresis corresponding to a previously reported autonomously replicating sequence (ARS), CARS2. We show that all the identified chromosomal origins are bound by *C. albicans* homologs of conserved pre-replication complex proteins Orc2 and Mcm2. Previous reports from our lab and others have shown that there is a strong inter-relationship between centromere function and pericentric origin activity, suggesting that these origins are epigenetically regulated. However, we find that short intergenic regions corresponding to each of these origins functions as an ARS element in circular plasmids. Further, we use the novel strategy of *in vivo* gap repair to demonstrate that circular ARS plasmids can exist independently *in vivo* in *C. albicans.* Taken together, these results show that DNA sequence underlies the function of both centromere proximal and centromere distal chromosomal origins in *C. albicans*.

## Introduction

The replication of a eukaryotic chromosome initiates from multiple distinct sites, called the replication origins, which ‘fire’ only once per cell cycle. The origin recognition (ORC) complex, a multi-subunit evolutionarily conserved protein complex, binds to these sites in an ATP dependent manner and marks the potential sites for origin assembly (Bell and Stillman, 1992). The Mcm or the minichromosome maintenance group of proteins have diverse functions ranging from replication to chromosome condensation (Forsburg, 2004). A group of 6 proteins, Mcm 2-7, form a ring shaped structure (Adachi et al., 1997) that is loaded at the G1 phase to origin sequences that are already bound by the ORC. Binding of the Mcm complex leads to the formation of the pre-replication (pre-RC) complex and licenses the origins for firing in the subsequent S phase (Chong et al., 1995). During S phase the Mcm complex acts as a replicative helicase that helps in unwinding of double stranded DNA ahead of the moving replication fork (Bell and Dutta, 2002).

In bacteria, the origin of replication (ori) is found to comprise of a common consensus sequence called the DnaA box, which is the binding site for the initiator protein DnaA (Messer and Weigel, 1997; Mott and Berger, 2007). In contrast, the initiation of DNA replication can occur on almost any genomic DNA fragment in most metazoa (Vashee et al., 2003). Archaeal and yeast origins of DNA replication occupy an intermediate ground. Their functionality depends on conserved sequence hallmarks such as ARS elements as well as genomic context along with structural cues (Leonard and Mechali, 2013). ARS elements were initially identified in *Saccharomyces cerevisiae* as genomic DNA elements that were able to confer replication ability to origin-less plasmids thus enabling them to transform host cells at a high frequency (Struhl et al., 1979). Sequence analysis and mutagenesis experiments identified a 11 bp AT-rich consensus sequence (WTTTAYRTTTW), the ARS consensus sequence (ACS), that was found to be essential for the ARS activity (Marahrens and Stillman, 1992). Subsequently, ARS elements have been identified in other budding yeast species including *Kluveromyces lactis, Yarrowia lipolytica* and fission yeast *Schizosaccharomyces pombe* (Maundrell et al., 1988; Irene et al., 2007; Vernis et al., 1997).

*Candida albicans* is a diploid human pathogenic budding yeast belonging to the CTG clade of hemiascomycota (Fitzpatrick et al., 2006; Legrand et al., 2019) Despite having superficial similarities to *S. cerevisiae*, the life cycle and genome organization of *C. albicans* reflects several unique features such as a regional epigenetically regulated centromere with unique DNA sequences (Baum et al., 2006; Sanyal et al., 2004), unusually long telomere repeats (McEachern and Hicks, 1993) and evidence of a parasexual cycle with a program of concerted chromosome loss (Bennett and Johnson, 2003; Legrand et al., 2019) However, very little is known about the basic biological process of replication in this organism. The study of replication origins in *C. albicans* had been initiated by testing the ARS property of plasmids carrying genomic DNA fragments in *S. cerevisiae* (Cannon et al., 1990; Herreros et al., 1992; Kurtz et al., 1987). The plasmids carrying these ARS elements were capable of autonomous replication and exhibited a high frequency of transformation in both *S. cerevisiae* and *C. albicans*. However, none of these ARS elements were tested for their chromosomal origin activity. In *S. cerevisiae* (Dubey et al., 1991) and *S. pombe* (Okuno et al., 1997), most, but not all, ARS elements corresponded to genomic origins. Therefore, several other methods such as genome-wide ChIP (chromatin immunoprecipitation)-on-chip (using antibodies against the conserved proteins of the pre-replication complex) (Heichinger et al., 2006; Wyrick et al., 2001) and two dimensional agarose gel electrophoresis assays (Brewer and Fangman, 1987) have been used either singly or in combination to study chromosomal origins of replication in these organisms. A study using the above techniques revealed conserved properties of *C. albicans* replication origins such as binding of ORC, nucleosome depletion and a primary sequence motif (Tsai et al., 2014). Moreover, the study concluded that there are two classes of replication origins, namely, the arm origins that are dependent on the primary sequence motif and the pericentric origins that are epigenetically regulated. In a parallel study, we had identified chromosomal origins flanking the centromeres of chromosome 5 and chromosome 7 in *C. albicans* using the 2-D agarose gel electrophoresis origin mapping technique (Mitra et al., 2014). In this paper, we have identified another non-centromeric origin at the locus of a known ARS element. We show that these chromosomal origins of DNA replication are bound by conserved members of the pre-RC complex, Orc2 and Mcm2. Further, we demonstrate that all the three tested origins show high transformation frequency and can exist as free circular plasmids in *C. albicans* cells, thus indicating that DNA sequence contributes to their function, irrespective of their genomic location.

## Results

### Identification of a centromere-distal chromosomal origin of replication in *C. albicans*

The first genome-wide replication study in *C. albicans* involved microarray analysis to ascertain the replication timing of the centromeres (Koren et al., 2010). Each centromere was found to be associated with the earliest firing origin in the genome and the presence of one such origin was verified by the 2-D gel analysis. Subsequently, while analyzing replication intermediates across the centromere of chromosome 7 (CEN7) by 2-D gel assays we had identified and reported four replication origins flanking CEN7 (pCENORI7-LI and pCENORI7-RI) and CEN5 (pCENORI5-LI and pCENORI5-RI) (Mitra et al., 2014). In this study, we initially tested whether a previously established *C. albicans* ARS sequence, CARS2 (Cannon et al., 1990), behaved as a chromosomal origin. BLAST alignment mapped CARS2 to an intergenic region flanking the RDN18 locus in chromosome R (Figure 1A). The analysis of replication intermediates of an EcoRI fragment containing CARS2 by the 2-D gel electrophoresis assay showed the presence of both Y arc and bubble arc signals (Figure 1A), indicating that this fragment contains an internal origin lying within its central third part. However, this origin, like the previously mapped origins (Mitra et al., 2014) is active only in a fraction of cells in the population and in other cells the origin containing fragment is passively replicated by forks coming from external origin(s). As all the origin-containing fragments produce prominent Y arc signals, none of these origins seem to fire as efficiently as some of the strong origins of *S. cerevisiae.* We named the newly found centromere distal origin as dCENORIR-RI. Thus, CARS2 is active as an origin of replication in its chromosomal locus in *C. albicans*.

**Figure 1.**
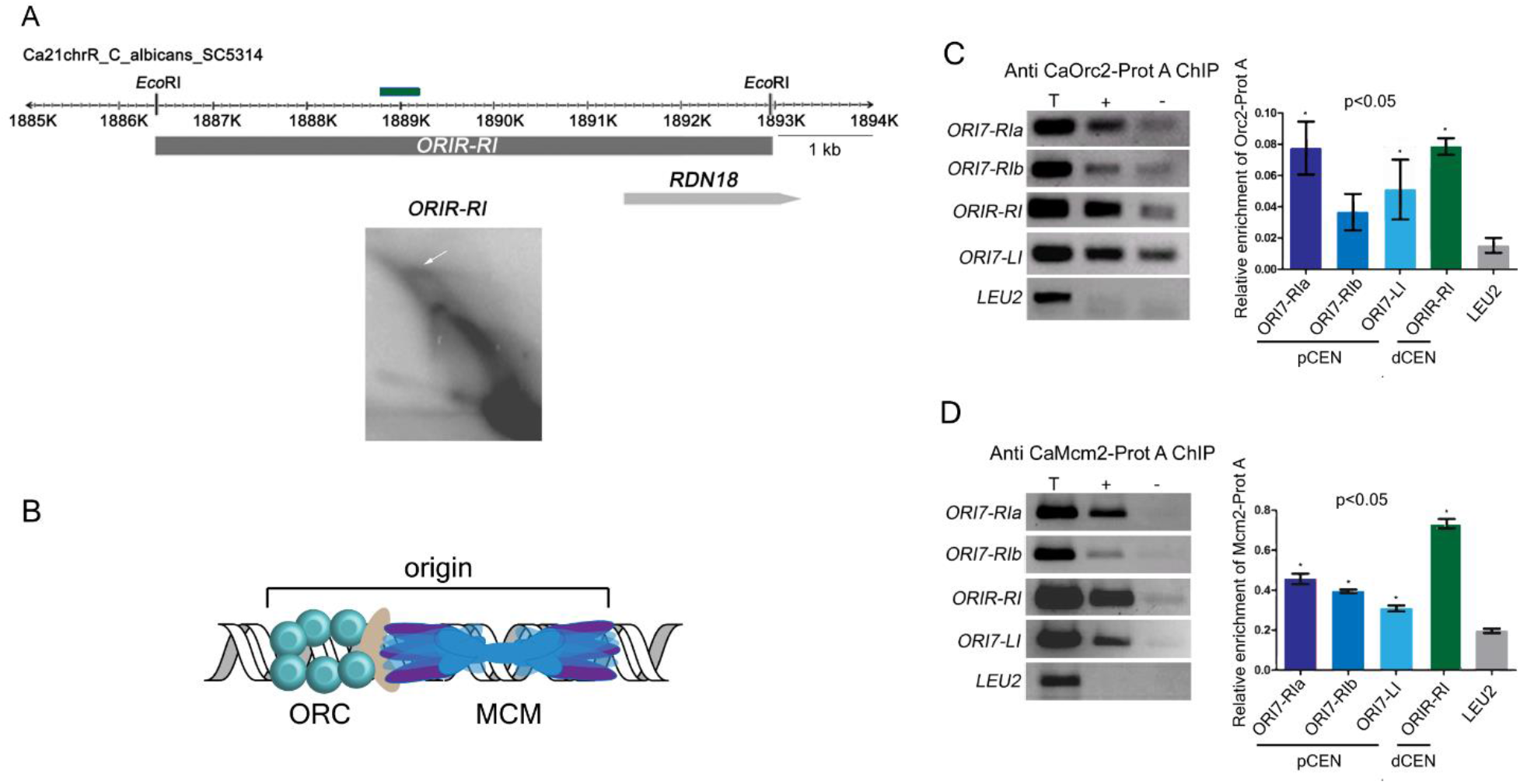
Centromeric (ORI7-RI, ORI7-LI) and non-centromeric (ORIR-RI) chromosomal replication origins identified by 2-D agarose gel electrophoresis in *C. albicans* are bound by CaOrc2 and Ca Mcm2. **(A)** A line diagram shows a ~10 kb region on chromosome R shows the position of ORIR-RI (filled grey rectangle). Arrowheads and numbers indicate the positions and the identities of the ORF. Coloured rectangles indicate qPCR primer position. 2D image shows simple ‘Y’ arc as well as the typical ‘bubble’ arc that indicates the presence of an active replication origin within the fragment. **(B)** Schematic showing the pre-RC consisting of ORC and MCM complexes bound to an origin. **(C)** Standard ChIP assays with anti-Prot A antibody followed by PCR (left) and quantitative real time PCR (qPCR) (right) were performed for quantifying the enrichment of CaOrc2-Prot A and **(D)** CaMcm2-Prot A at ORI7-RI, ORI7-LI and ORIR-RI. Amplification from a control gene region (LEU2) was also performed to detect the background DNA elution in the ChIP assays. Enrichment of ORC and MCM proteins at each specified location was calculated as relative enrichment and values were plotted as mean of three technical replicates ± SD.

Overlapping binding of two distinct sub-complexes of the pre-replication complex proteins (Figure 1B) have been used as a standard method of identifying genome-wide replication origins by ChIP-chip or ChIP-sequencing assay. Towards that objective, we epitope tagged one allele of the MCM and ORC genes with a C terminal Tandem Affinity Purification or TAP epitope (Calmodulin Binding Domain – TEV-Protein A) in a wild-type *C. albicans* strain BWP17 resulting in CAKS107 (MCM2/MCM2-TAP) and CAKS108 (ORC2/ORC2-TAP). Using these strains, we looked at the enrichment of Orc2 and Mcm2 at the 2-D identified chromosomal origins. ChIP assays were performed in asynchronous cells with anti-Protein A antibodies for CaOrc2-Prot A and CaMcm2-Prot A. Immunoprecipitated DNA was PCR amplified by primers corresponding to dCENORIR-RI, pCENORI7-RI and pCENORI7-LI. ChIP-PCR analysis confirmed the enrichment of CaOrc2 and CaMcm2 at pCENORIR-RI, pCENORI7-LI and dCENORI7-RI in the ‘+’ antibody sample as compared to the ‘-’ or ‘beads-only’ control (Figure 1C and 1D (left)). In contrast, these MCM and ORC proteins were not enriched on a genic region (*LEU2*). We further verified our results by performing quantitative real-time PCR (qPCR) reactions for the same regions (Figure 1C and 1D (right)). All the three regions showed significant enrichment of CaOrc2 and CaMcm2, as compared to the control *LEU2* locus. Thus, we validated that the pre-RC complex is indeed loaded on to the chromosomal origins identified by the 2D gel assay.

### DNA sequence contributes to origin function in *C. albicans* irrespective of their chromosomal location

In budding yeasts all chromosomal origins have been found to have ARS activity (Huberman et al., 1988) with the ARS sequence always localized to an intergenic region (Brewer, 1994). We had previously showed that intergenic fragments from pCENORI7-LI and pCENORI7-RI show ARS activity (Mitra et al., 2014). However, we had not determined the transformation efficiency of these plasmids nor was it determined whether these plasmids are maintained in free form *in vivo*. The intergenic regions from pCENORI7-RI and pCENORI7-LI were named as CARS4 and CARS5 respectively. The corresponding plasmids were renamed as pARS4-1 and pARS5 (Figure 2A and S1A). Tandem copies of ARS sequences in a plasmid have been shown to increase the ARS efficiency (Hyman and Garcia-Garcia, 1993). Hence the plasmid pARS4-3 was constructed, having two copies of CARS4 (Figure 2A). Transformation of these plasmids into *C. albicans* strain BWP17 (Baum et al., 2006) yielded a high frequency of transformation (ranging from 5 x10^2^ to 2 x10^3^ transformants per μg of plasmid DNA transformed) (Figure 2B and S1B). These values were comparable with the range of transformation frequency described for previous *C. albicans* ARS elements (2 x 10^2^ to 1 x10^4^ transformants per μg of plasmid DNA transformed) (Cannon et al., 1990; Herreros et al., 1992). We further cloned CARS2 into the same backbone as pARS4 to get the plasmid pARS2 (Figure 2A). Transformation of this plasmid into *C. albicans* yielded comparable results as previously obtained (Cannon et al., 1990), re-confirming that pARS2 functions as an ARS plasmid (Figure 2B). As a negative control, we cloned a random ~2.5 kb genomic DNA fragment into the pUC19-URA3 backbone to form the plasmid pCtrl-1. Transformation of this plasmid produced only a few large, smooth and round colonies indicative of integrants (Figure S1C).

**Figure 2.**
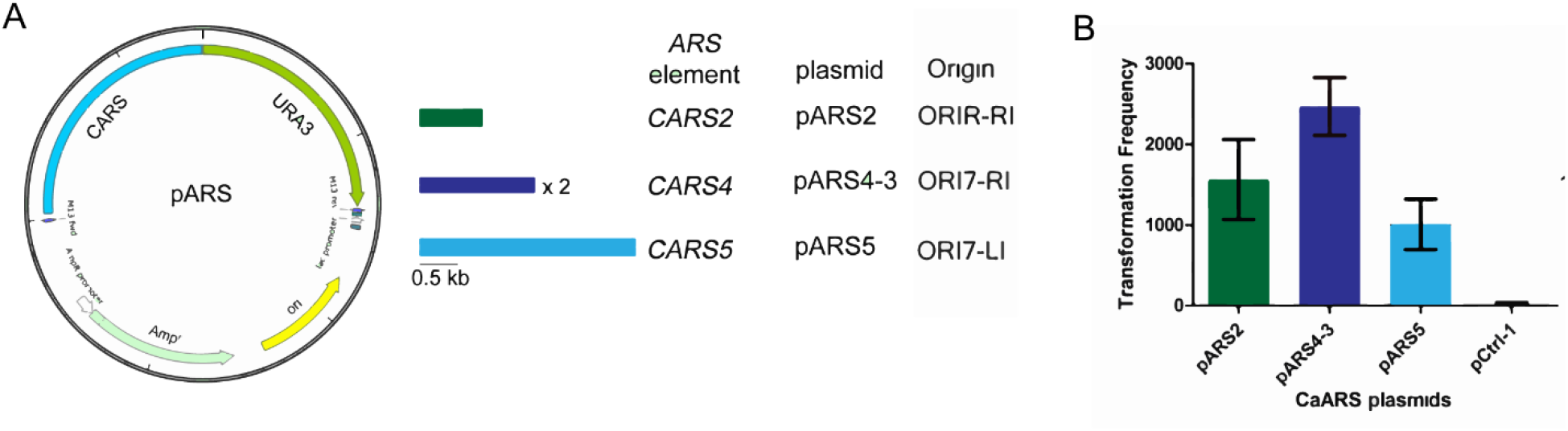
Chromosomal origins are associated with ARS activity in *C. albicans*. **(A)** The plasmid map of the CARS backbone, the relevant ARS fragments and their corresponding chromosomal origins, and plasmid names are shown. **(B)** Bar graphs show the transformation frequency (number of transformants /μg of transforming DNA) of the different classes of CARS plasmids. Each bar indicates the mean transformation frequency of three independent transformation experiments ± SD.

In order to know the *in vivo* status of these ARS plasmids, we isolated total genomic DNA from *C. albicans* ARS transformants and performed Southern hybridization with a sequence probe taken from the backbone of the vector E. coli pUC19 (Figure 3A). Three transformants were analyzed for each of pARS2, pARS4-3 and pARS5. The untransformed wild-type genomic DNA as well as the free pARS2, pARS4-3 and pARS5 plasmids isolated from *E. coli* were taken as controls. The results showed that at least one transformant from each class of ARS plasmids showed the presence of free monomeric plasmid (denoted as * in Figure 3B). Apart from that, signals were observed at high molecular weight positions which indicated the occurrence of genomic integration or plasmid multimers (denoted as ** in Figure 3B). Such oligomeric formations had been reported earlier for *C. albicans* ARS plasmids (Kurtz et al., 1987). Thus, it was observed that circular *C. albicans* ARS plasmids exist as a mixture of free monomers and genomic integrants/ concatenated multimers *in vivo*.

**Figure 3.**
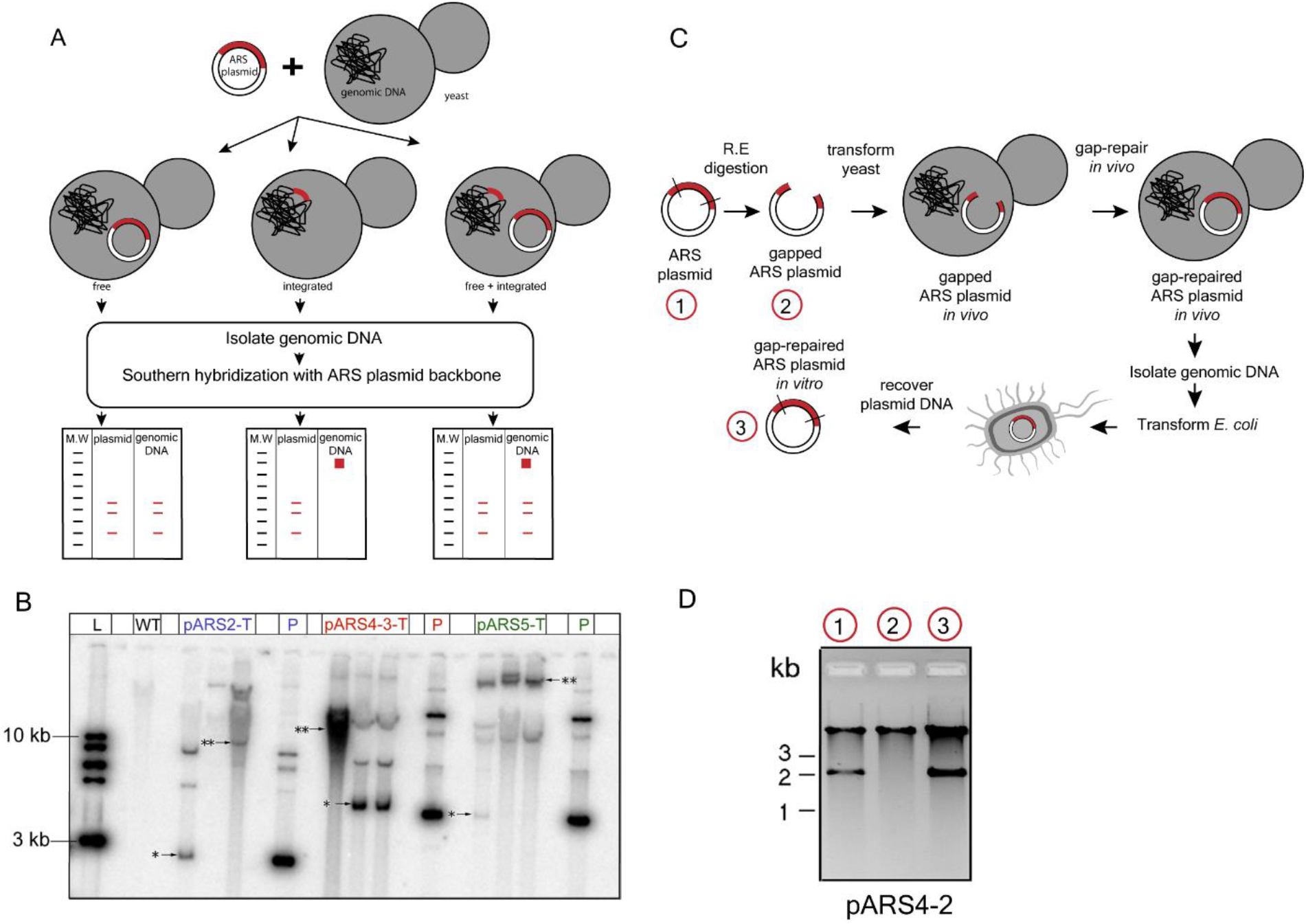
Chromosomal origins can exist as free extrachromosomal circular plasmids *in vivo.* **(A)** Schematic showing the strategy for the detection of ARS plasmid states *in vivo.* Briefly genomic DNA isolated from *C. albicans* cells transformed with ARS plasmids were probed by Southern hybridization using the plasmid backbone to detect ARS plasmid states *in vivo* **(B)** Phosphorimager picture of Southern blots of *C. albicans* CARS transformants (pARS2-T, pARS4-3-T and pARS5-T) hybridized by a probe obtained from the backbone sequence of E. coli present in the plasmid. Untransformed wild-type DNA and the respective free plasmids (denoted as P) are taken as controls. 1 kb ladder is used as a molecular weight marker. * indicates the position of plasmid monomer and ** indicates the position of higher molecular weight multimers. **(C)** A strategy of *in vivo* gap-repair of a Ca-ARS plasmid in order to demonstrate the *in cellulo* existence of independent circular ARS. **(D)** An example of repair of gapped pARS4-2 plasmid. Plasmid DNAs were isolated from E. coli, digested with EcoRV and run on a 0.8% agarose gel. Lane 1, pARS4-2; lane 2, gapped and recircularized pARS4-2; lane 3, gap-repaired pARS4-2. Presence of the 2 kb band in lane 3 indicates the *in vivo* repair of the gap in pARS4-2.

We addressed the question of *in vivo* status of a circular *C. albicans* ARS plasmid in a second way by taking a novel approach of *in vivo* gap repairing an ARS plasmid and subsequently testing whether the free plasmid can be recovered in *E. coli* (Figure 3C). Gap repair is a homology mediated recombination process which repairs a double stranded gap in a plasmid using a chromosomal repair template (Orr Weaver and Szostak, 1983). Gap repair has never been shown to occur in *C. albicans.* In order to clone CARS4 by gap repair in a plasmid, we digested the pARS4-2 plasmid (Figure 3D) with EcoRV to create a 2 kb gap including the CARS4 region. This gapped plasmid was transformed into *C. albicans.* Total genomic DNA isolated from the transformants was then transformed into *E. coli* in order to recover gap-repaired plasmids, if any. We were able to observe repair of a gap in at least two independent *C. albicans* transformants (Figure 3D). This result not only confirmed that pARS4-2 was able to exist independently *in vivo* but also demonstrated that, *in vivo* gap repair occurs and can be used as a tool in *C. albicans*.

## Discussion

In the present study we have demonstrated that a known *C. albicans* ARS plasmid behaves as an origin in its native chromosomal locus. In all, we have characterized three chromosomal origins, two of them pericentric (Mitra et al., 2014) and a third one in the chromosomal arm (present study). We confirmed the functionality of these origins by showing that they are bound by native *C. albicans* pre-RC proteins Orc2 and Mcm2.

In recent years novel non-canonical roles have been identified for both ORC and MCM proteins. In humans Orc2 was found to be localized at CEN in a HP1 mediated manner and helped in pericentric heterochromatin organization (Prasanth et al., 2005, 2010). Recently it was shown that Mcm2 acts as a chaperone for parental H3 (Huang et al., 2015) and CENP-A (Zasadzińska et al., 2018) during replication. Orc4 was also found to be important for centromere function in *C. albicans* (Sreekumar et al., 2018). An earlier study (Koren et al., 2010) in *C. albicans* had indicated that the function of replication origins flanking the centromeres are governed by centromere function. Subsequently it was found in *S. cerevisiae* that kinetochores recruit Dbf4, thereby regulating timing of firing of centromere flanking origins (Natsume et al., 2013). Further, we found that deletion of a centromere flanking replication origin reduces centromere function. Recently it has been suggested that a ‘replication wave’ propagates from early replicating centromeric origins to the late replicating telomeric origins (Lazar-Stefanita et al., 2017). All these studies seem to indicate that centromere flanking origins are epigenetically regulated.

In order to determine whether the underlying DNA sequence plays any role in the function of the replication origins that we identified, we tested them for ARS activity. A previous study (Tsai et al., 2014) had suggested that circular *C. albicans* ARS plasmids get integrated into the genome and hence are not suitable for studying sequence requirements of origin function. In our study, we obtained a clear difference in transformation frequency (800-2500 transformants per μg of transforming DNA) of circular ARS plasmids as compared to a random control plasmid (~30 transformants per μg of transforming DNA). Moreover, a greater proportion of the circular ARS transformant colonies was small and wrinkled, which is a characteristic feature of free ARS plasmids that are mitotically unstable and lost at a high rate. In the same context, majority of the transformants on the control plate were large and round, indicating their stable integration into the genome. Finally, we proved by two approaches the extrachromosomal existence of circular ARS plasmids at the DNA level. The free circular monomer could be identified by radioactive probing after separation from the *C. albicans* genomic DNA on an agarose gel. We used a novel approach of gap repairing a circular ARS plasmid *in vivo*.

Gap repair, also known as gap-filling or *in vivo* cloning is a widely used technique in *S. cerevisiae* for error-free construction of plasmids within yeast cells that can restore gapped plasmids using homologous sequences as templates. Although the homologous recombination pathway is conserved in *C. albicans*, the occurrence of gap-repair has never been tested in this organism. Using restriction enzymes, we excised a fragment out of a Ca-ARS plasmid in order to generate a gapped template. Transformation of *C. albicans* cells with this plasmid and subsequent recovery of gap-repaired plasmids from *E. coli* demonstrated that 1) gap-repair occurs in *C. albicans* and 2) ARS plasmids can exist as independent extrachromosomal circles *in vivo*. Therefore, our present study shows, even for centromere flanking origins, the underlying sequence contributes significantly towards origin function, as, the sequence alone, when taken out of chromosomal context into a plasmid, shows ARS function. In *S. cerevisiae* origin function is strictly governed by the underlying sequence (ACS element). Despite that, it has been observed that surrounding chromatin landmarks such as centromeres and telomeres influence timing of DNA replication origin firing. We predict a similar situation in *C. albicans*, whereby, all origin functions are governed genetically by the DNA locus. However, the time of firing is modulated by the surrounding chromatin atmosphere. Since gap-repair cloning is not limited by the length of the insert to be cloned, our future objective will be to use this strategy to restore larger stretches of genomic DNA surrounding chromosomal origins in order to recapitulate the chromatin environment of origin activity.

## Abbreviations

ARS: autonomously replicating sequence
Orc2: Origin Recognition Complex Subunit 2
Mcm2: minichromosome maintenance protein 2
ORC: origin recognition complex
pre-RC: pre-replication complex
ACS: ARS consensus sequence.

## Acknowledgements

The work was supported by a grant from the Department of Biotechnology, Government of India to KS and DDD (BT/PR13724/BRB/10/782/2010).The intramural support from JNCASR is also acknowledged. SM was supported by a Council for Scientific and Industrial Research (CSIR), India senior research fellowship and a research associate fellowship from JNCASR.

## Materials and Methods

### Strains, primers, media and growth conditions

*C. albicans* strain BWP17 (Fonzi and Irwin, 1993) was grown in YPDU (1% yeast extract 2% peptone 2%dextrose 0.010% uridine) or SD (0.67% yeast nitrogen base, 2% dextrose) supplemented with amino acids (0.01%) as necessary at 3Ü°C, as described previously (Sanyal and Carbon, 2002). E. coli strain DH5α was used for all bacterial transformations. The *C. albicans* strains used in this study have been listed in Table S1. The primers used in this study have been listed in Table S2.

### 2-D agarose gel electrophoresis

Neutral-neutral (N-N) 2-D agarose gel electrophoresis of DNA replication intermediates was performed as described previously (Friedman and Brewer, 1995). Briefly, high quality genomic DNA was extracted from log-phase *C. albicans* cells by the CsCl density centrifugation method. Benzoylated Napthoylated DEAE (BND) cellulose fractionation was performed on this DNA to enrich for single stranded DNA. The enriched DNA was loaded onto 0.4% agarose gel in 1X TBE (Tris-HCL, Boric acid, EDTA) buffer and run for 15-16 hours at 15-20 volts at 4°C. The DNA molecules were separated according to their mass in the first dimension. After first dimension, the desired lanes were cut and run in the second dimension (at 90° to the first) in 1.1% agarose gel in 1X TBE for 2-4 hours at 100-125 volts in presence of ethidium bromide (0.3 mg/ml) at 4°C. The high voltage and higher percentage of the second dimension gel separated the linear DNA from branched DNA molecules. Southern transfer was done under alkaline transfer conditions. The blot was hybridized with probes from the upstream and downstream regions of centromere 7 (CEN7) and from rDNA locus of chromosome R. Hybridized membranes were exposed to Phosphorimager films and images captured were analyzed by Phosphorimager using the Image Reader FLA5000 Ver.2 software (Fujifilm).

### Construction of plasmids

For constructing *C. albicans* ARS plasmids the CaURA3 gene was digested as a 1.3 kb HindIII fragment from the plasmid pCaDis (Care et al., 1999) and cloned into the HindIII site of pUC19 to get pUC19-URA3. Subsequently a 5 kb region downstream of CEN7 showing origin signal in 2-D gel assay was amplified from *C. albicans* genome and cloned as a SmaI (S) fragment into pUC19-URA3, to get pARS4-2. A 1.4 kb intergenic region within the 5 kb fragment was separately cloned into the pUC19-URA3 as SmaI fragments, both in one and two copies, thus generating the plasmids pARS4-1 and pARS4-3 respectively. The previously reported *C. albicans* ARS, CARS2 (Cannon et al., 1990), was amplified from the plasmid pAB1 (Baum et al., 2006) and cloned as a SmaI fragment into the same backbone to get pARS2.

### Construction of *C. albicans* strains

CAKS107 (MCM2/MCM2-TAP) was constructed by integrating a C-terminal Prot A tagging cassette with NAT marker created by overlap extension PCR using the primers M1 to M6 (Table S2) in BWP17 (Table S1). Integration of the cassette at the desired site was confirmed by PCR using the primers Mcm2cf1 and TAP-Conf RP (Table S2).

The upstream fragment (384 bp) and the downstream fragment (615 bp) of CaORC2 were PCR amplified from the *C. albicans* genome with the primer pairs CAO2-TAP-1,2 and 3,4 respectively (Table S2) and were cloned in the XbaI / BamHI and ApaI / KpnI sites of pPK335 plasmid (TAP-URA3) (Corvey et al., 2005). The 2965 bp XbaI / KpnI fragment was gel extracted and transformed into BWP17 cells (Table S1) and the transformants were confirmed by PCR with primer pair CAO2-TAP-CFP and TAP-RP (Table S2) thus generating CAKS108 (ORC2/ORC2-TAP) strain (Table S1).

### *C. albicans* transformation

Transformation of *C. albicans* was performed using the lithium acetate mediated transformation technique, as described previously (Baum et al., 2006). Transformation was also performed by a modification of the spheroplasting method described by Burgers and Percival (Burgers and Percival, 1987). A 50 ml culture of *C. albicans* was grown in YPD (U) medium till an O.D600 of 1.2 and the cells were harvested by centrifugation (5 min at 4000 rpm). The cells were washed with 20 ml of autoclaved distilled water and 20 ml of 1M sorbitol successively. Then the cells were resuspended in 20 ml of 1.2 M sorbitol, 20 mM Na-HEPES (pH 7.4) to which 2 μl of β-mercaptoethanol was added. Approximately 90 μl of lyticase (10 mg/ ml) was added and the cells were incubated at 30°C with 65 rpm for spheroplasting. O.D800 was measured every 15 min to determine the percentage of spheroplasting and the cells were harvested (5 min at 2000 rpm) when cells were 90% spheroplasted The spheroplasts were resuspended in 20 ml of 1M sorbitol with gentle shaking and then harvested by centrifugation (5 min at 2000 rpm). The above step was repeated with 20 ml of STC buffer (1M sorbitol, 10 mM Tris pH 7.5, 10 mM CaCl2). Finally, the spheroplasts were resuspended in 1 ml of STC buffer. Transforming DNA was added to 0.1 ml aliquot of the spheroplast suspension, along with 1 μl of single stranded carrier DNA (10 mg/ml) and incubated at room temperature for 10 min. Subsequently, 1 ml of PEG solution (10mM Tris pH 7.5, 10 mM CaCl2, 20% PEG 8000) was added to the above suspension and mixed gently. After a further incubation at room temperature for 10 min, the suspension was spun down at 2000 rpm for 5 min and resuspended in 150 μl of SOS buffer (1 M sorbitol, 6.5 mM CaCl2, 0.25% yeast extract, 0.5% bactopeptone) for physiological recovery. The spheroplast suspension was incubated at 30°C for 45 minutes. About 8 ml of TOP agar (1M sorbitol, 2% agar, 0.67% nitrogen base, 3% glucose) (kept at 46-50°C) was added to the suspension, mixed by inversion and immediately poured directly onto the surface of selective media and incubated at 30°C.

### ARS activity assays

*C. albicans* cells were transformed by the spheroplast method with ARS plasmids and the number of colonies on selective plates was generally scored for up to 1 week. The ARS activity of each plasmid was expressed as the transformation frequency (number of transformants /μg DNA). Each transformation frequency is an average of three independent transformation experiments. Error bars indicate standard deviation (S.D) from the mean.

### Chromatin Immunoprecipitation (ChIP)

ChIP-PCR analysis was done as described previously (Sanyal et al., 2004). Rabbit anti-Protein A antibodies (Sigma Cat. No P3775) was used for ChIP at a final concentration of 20 μg/ml of immunoprecipitate (IP). Asynchronous cultures of *C. albicans* strains were grown in YPDU till O.D600 of 1.0. They were cross-linked with 37% formaldehyde for 1 h. Subsequently, sonication was performed with Biorupter (Diagenode) to get sheared chromatin fragments of an average size of 300-500 bp. The fragments were immunoprecipitated with anti-Protein A antibodies (Sigma). ChIP DNA was analyzed by both semi-quantitative PCR and quantitative real-time PCR (qPCR) using primer pairs that amplify selected regions from ORI7-RI, ORI7-LI and ORIR-RI. Amplification from a non-centromeric control region (LEU2) was also performed to detect the background immunoprecipitated DNA. For anti-Protein A ChIP analysis, qPCR was performed on a Rotor Gene 6000 real-time PCR machine with IQ Sybr Green Supermix (Bio-Rad). Cycling parameters were as follows: 94°C/30 s, 55°C/30 s, 72°C/45 s repeated 40X. Melt curve analysis was performed from 55°C to 94°C. Error bars were calculated as S.D. for three technical replicates of each ChIP sample.

### ChIP-qPCR enrichment calculation

The ChIP-qPCR enrichment was determined by the fold enrichment method. In brief, the Ct values for input were corrected for the dilution factor and then the percent of the input chromatin immunoprecipitated by the antibody was calculated as Ct+Ab/Adjusted CtInput. This was subtracted from Ct-Ab/Adjusted CtInput to generate the fold enrichment values. One way ANOVA and Dunnett post tests were performed to determine statistical significance.

### *E. coli* transformation

*E. coli* transformations were performed by standard chemical and electroporation methods.

