## Supplemental data for "Chromosomal replication origins in *Candida albicans* are genetically defined irrespective of their chromosomal context"

Figure S1

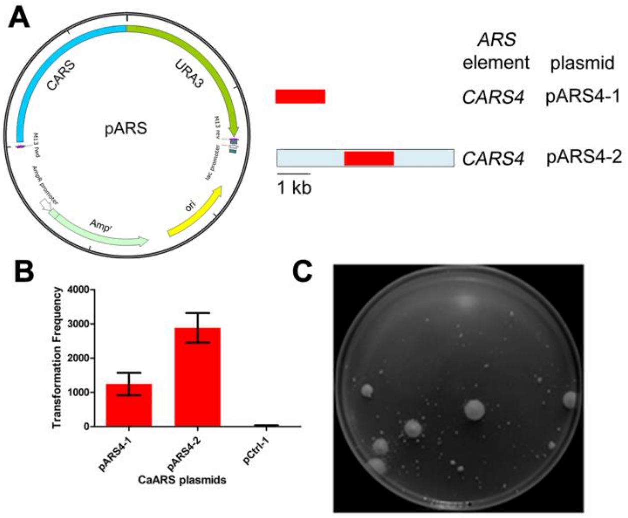


**Figure S1. Chromosomal origins are associated with *ARS* activity in *C. albicans*.** (A) The plasmid map of the *CARS* backbone, the relevant *ARS* fragments and plasmid names are shown. pARS4-2 contains the entire ~5 kb EcoRI fragment (*ORI7-RI*) that detected origin activity in the 2-D analysis in Fig. 3. **(B)** Bar graphs show the transformation frequency (number of transformants /µg of transforming DNA) of the different classes of *CARS* plasmids. Each bar indicates the mean transformation frequency of three independent transformation experiments ± SD. **(C)** The plate picture shows the results of an *ARS* function assay using the control plasmid pCtrl-1 (containing a random ~1.4 kb intergenic fragment).

**Table S1.** Strains used in this study.

| Strain | Genotype | Reference |
|---|---|---|
| BWP17 | *Δura3::imm434/Δura3::imm434, Δhis1::hisG/Δhis1::hisG, Δarg4::hisG/Δarg4::hisG* | (Wilson, et al. 1999) |
| CAKS107 | *Δura3::imm434/Δura3::imm434, Δhis1::hisG/Δhis1::hisG, Δarg4::hisG/Δarg4::hisG, MCM2/MCM2-TAP(NAT)* | This study |
| CAKS108 | *Δura3::imm434/Δura3::imm434, Δhis1::hisG/Δhis1::hisG, Δarg4::hisG/Δarg4::hisG, ORC2/ORC2-TAP(URA3)* | This study |
| Plasmid | Description | Reference |
| pKS101 | pUC19 + CaURA3 | (Mitra, et al. 2014) |
| pARS4-1 | pKS101 + 1.4 kb intergenic region from *ORI7-RI* (1 copy) | (Mitra, et al. 2014) |
| pARS4-2 | pKS101 + 5 kb *ORI7-RI* | This study |
| pARS4-3 | pKS101 + 1.4 kb intergenic region from *ORI7-RI* (2 copies) | This study |
| pARS5 | pKS101 + 2.7 kb intergenic region from *ORI7-L1* | (Mitra, et al. 2014) |
| pCtrl-1 | pKS101 + 1.2 kb random intergenic region | This study |

Table S2. Primers used in this study.

| **Primers** | **Sequence** (5' to 3') |
|---|---|
| 7DS1F | CAATGGAACGGTTATCACTT |
| 7DS1R | AGCTGGTTTGTGAGTTAGGA |
| 7DS3F | AAAGAGCAGTTTCAGATCCA |
| 7DS3R | TCAACCGGATATTGTCTACC |
| FARS2 | TCCCCCGGGAAAAATCCGGGGTAGCGATG |
| RARS2 | TCCCCCGGGTGCAGCTACGAATGTTAGAG |
| 7ori 1a | TCCCCCGGGCGTCTGTTTCAGAGTCTGG |
| 7ori 1b | TCCCCCGGGGTCTCGATCTCACCAATAG |
| 7ori 2a | TCCCCCGGGTGTCACAGGCATTGTGTAG |
| 7ori 2b | TCCCCCGGGAGCTGGTTTGTGAGTTAGG |
| M1 | AATTTTATCCAGATTTGATATTATG |
| M2 | TTCCATCTTCTCTTTTCCATCAAAGTATATTTCATAAATTTACTTC |
| M3 | GAAGTAAATTTATGAAATATACTTTGATGGAAAAGAGAAGATGGAA |
| M4 | TACAAACAATAACAATAACTATAACGATATCAAGCTTGCCTCGTC |

| | |
|---|---|
| M5 | GACGAGGCAAGCTTGATATCGTTATAGTTATTGTTATTGTTTGTA |
| M6 | TTCAAGATATTATAAAATAGTCGAA |
| TAP-Conf RP | CGTTAGCGCTTTGGCTTGGGTCATC |
| Mcm2cf1 | AATCCTAATGGAGGTCGATA |
| CARS2-1 | ACAACCGGGATAGTGTTTGG |
| CARS2-2 | AATATGTATGTCCGGGTGGC |
| 2498-19 | GCCATACGGTAGTCAAACTCCTGG |
| 2498-20 | CCTGAACCACTACTGCAGAAACGT |
| CaCh7F3 | ACAGCCACAGTAGTTCCAAT |
| CaCh7R3 | TGAAATCCCAGAATGCCGTAAG |
| CaCh7F4 | TCTACCACCGTACTTTTGTTGG |
| CaCh7R4 | AATGACGTTTTGATGCATATTGC |
| 2498-20RTR | TTTAGTTGCCAGACCTTCGC |
| CaCh7F3RTF | CTGGTATTCACAATGGAACGGT |
| CaCh7F3RTR | GTCACCCCAATTCAAATCACGT |
| CaCh7F4RTF | GGAGCTGGCGATCAATTTGT |
| CaCh7F4RTR | TCACACATGAGAGGACCGTT |

| | |
|---|---|
| CAO2-TAP-1 | GCT CTA GAC GTG ATA AGT TTA GGC AAA TCC |
| CAO2-TAP-2 | CGG GAT CCT ACA TCA AAT TCT TGC TTA TAT ATG |
| CAO2-TAP-3 | AGG GGC CCG CTC AAC TAA TTT ACA TTT TCT CC |
| CAO2-TAP-4 | GGG GTA CCT GGC TCT GTT AGT TTT GTT TTG |
| CAO2-TAP-CFP | TCA CAA CAT ATG CCA CTT ACC AAC |
| TAP-RP | AGG TTA GCG CTT TGG CTT GGG TCA TC |
